# RootNet: A Convolutional Neural Networks for Complex Plant Root Phenotyping from High-Definition Datasets

**DOI:** 10.1101/2020.05.01.073270

**Authors:** Robail Yasrab, Michael P Pound, Andrew P French, Tony P Pridmore

**Affiliations:** Computer Vision Laboratory, School of Computer Science, The University of Nottingham, Nottingham, NG8 1BB, United Kingdom

**Keywords:** Convolutional Neural Network (CNN), Plant phenotyping, PyTorch, Residual Network, Roots, Computer Vision, Plants

## Abstract

Plant phenotyping using machine learning and computer vision approaches is a challenging task. Deep learning-based systems for plant phenotyping is more efficient for measuring different plant traits for diverse genetic discoveries compared to the traditional image-based phenotyping approaches. Plant biologists have recently demanded more reliable and accurate image-based phenotyping systems for assessing various features of plants and crops. The core of these image-based phenotyping systems is structural classification and features segmentation. Deep learning-based systems, however, have shown outstanding results in extracting very complicated features and structures of above-ground plants. Nevertheless, the below-ground part of the plant is usually more complicated to analyze due to its complex arrangement and distorted appearance. We proposed a deep convolutional neural networks (CNN) model named “RootNet” that detects and pixel-wise segments plant roots features. The feature of the proposed method is detection and segmentation of very thin (1-3 pixels wide roots). The proposed approach segment high definition images without significantly sacrificing pixel density, it leads to more accurate root type detection and segmentation results. It is hard to train CNNs with high definition images due to GPU memory limitations. The proposed patch-based CNN training setup makes use of the entire image (with maximum pixel desisity) to recognize and segment give root system efficiently. We have used wheat (Triticum aestivum L.) seedlings dataset, which consists of wheat roots grown in visible pouches. The proposed system segments are given root systems and save it to the Root System Markup Language (RSML) for future analysis. RootNet trained on the dataset mentioned above along with popular semantic segmentation architectures, and it achieved a benchmark accuracy.

## I. Introduction

Recently plant phenotyping is getting more attention due to the ever-growing human population. Plant genomics research helps to improve plant growth and productivity [1]. However, it requires a very comprehensive understanding of plant growth and cultivation patterns. Currently, machine learning and computer vision approaches play an important role in supporting image-based plant phenotyping. Singh et al. [2] proposed that complex plant phenotyping problems can be resolved using state-of-the-art machine learning techniques. There are two main areas that biologist focused on when studying plants and crop species, the above-ground (stem, leaves, branches, fruits, seeds, etc.) and below-ground (primary roots, lateral roots, storage roots, etc.) [3]. The most imagebased plant phenotyping has focused on the above-ground plant phenotyping (easy availability of image datasets), though the root phenotyping is not as popular. However, it is also necessary for determining plant growth. The reason for this is that below-ground plant phenotyping is an expensive and challenging task. Roots may usually grow in the soil and measuring their traits may sometimes be impossible without damaging and influencing their natural growth process. Therefore, root phenotyping is not getting that much attention as above-ground plant phenotyping got [4], [5], [6]. Classification and segmentation using Deep learning usually require large datasets with good quality images [2], [7], which thus makes roots unpopular for phenotyping plants. However, the recent success of deep learning and imaging tools for several industrial and research facilities triggered their use for plant phenotyping applications. The result is many large and well-annotated datasets are now available for plant and crop traits classification and segmentation.

Recently, several researchers turned their attention toward the below-ground plant phenotyping and analysis. Inspired by the work of pound et al. [7], [6], [8], [9], the proposed research focused on roots segmentation and analysis. The key objective of this research is to explore the roots network and segment its complicated features more elaborately. However, in root phenotyping, the biggest challenge is the availability of root features to detect and segment them into different root types correctly. According to Dodge et al. [10], image quality is essential to machine learning models. Higher-resolution images have more details and provide robust features for deep learning. The neural network model learns subtle characteristics between visually similar classes when trained with high-resolution images for fine-grained image segmentation tasks. However, low-resolution images used for the same task misses some feature details. In other words, they learn the global context from low-resolution images and local context from high-resolution images. Thus, there is a trade-off between the two types of images, which sometimes depends on the context in which we implement the deep learning systems. Therefore, like in classic computer vision problems, we have made use of multiple resolutions images, to take advantage of both global and local context. It is challenging to train CNN with higher resolution images due to the limitations of the Graphical Processing Unit (GPU)/ computer main memory. Training CNN with low-resolution images possibly leads to loss of root details (especially for lateral root (second-order roots) hairs) those are merely 1-3 pixels in width. We addressed this challenging problem of segmentatiing high definition plant roots by developing a deep convolutional neural network (CNN) to separate images of whole wheat roots systems from their background. The wheat roots system is composed of primary and lateral roots. Primary roots are usually easy to detect and segment, but lateral roots are not due to their small size and complicated shape. RootNet designed to detect and segments primary and lateral roots, which are usually undetectable by traditional image-based systems. This proposed system also offers an additional feature to save the Root System Architecture (RSA) to Root System Markup Language (RSML) [11]. The RSML is a globally accepted standard for analysis and annotation of the root systems. Several well know tools [12], [13], [11] support RMSL for roots features analysis. We used the wheat (Triticum aestivum L.) root dataset for training and testing the proposed model. The resultant network outperforms early available semantic segmenting architectures. A graphical introduction of the proposed RootNet system is shown in Figure 1.

**Figure 1:**
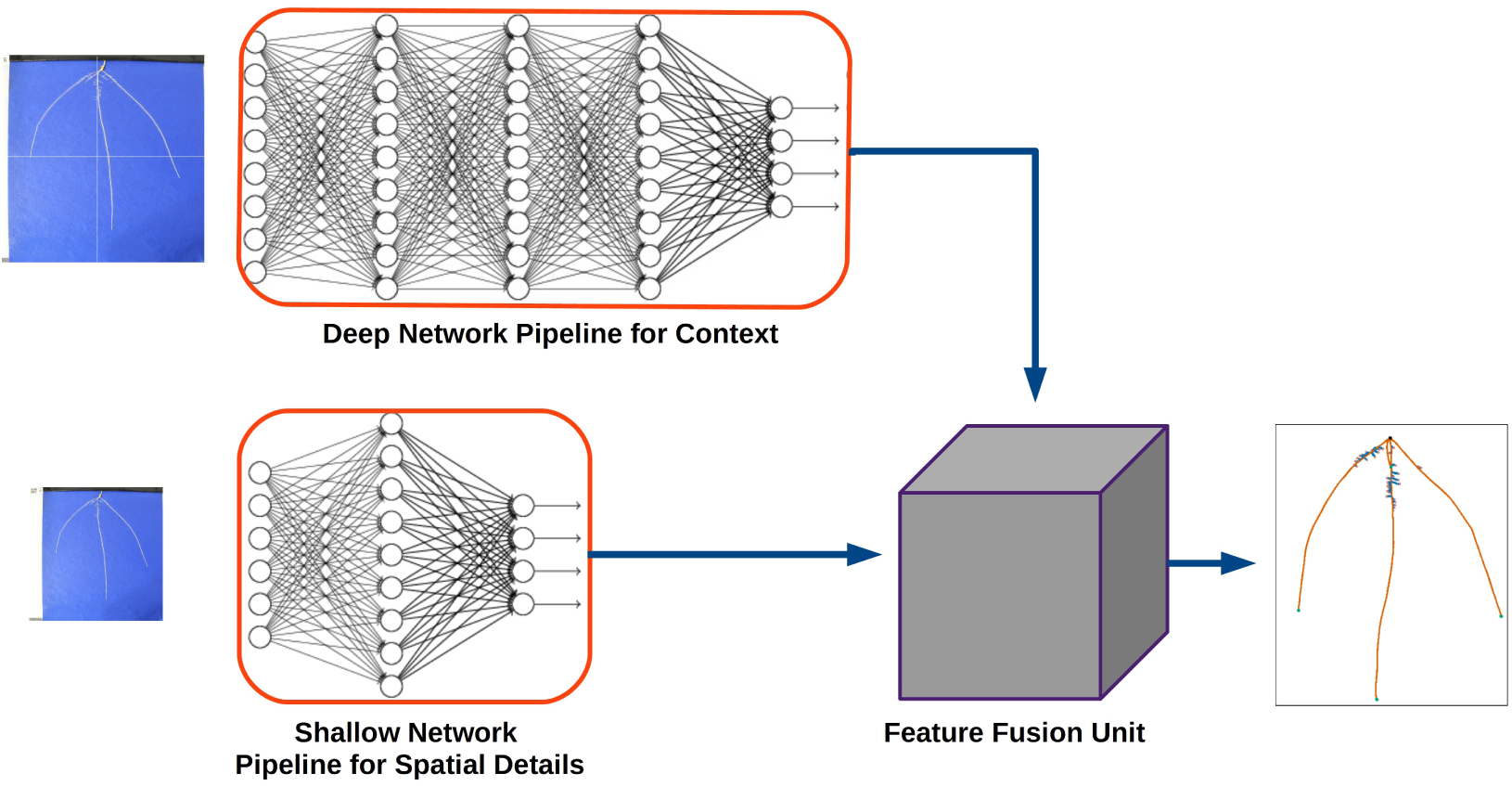
Basic Design of RootNet

The rest of the paper outlined as follows: The next section describes the background of plant phenotyping and its connection with machine learning. The following section is about the literature review, where we have described the earlier computer vision-based research contributions for plant phenotyping. The next section describes the methodology of the proposed network and design. Later, the section based on different experiments performed to assess CNN authenticity. The next part is about the results and analysis, and lastly, the conclusion and references section.

### A. Research Contributions

The main contribution of this research is that we have designed a state-of-the-art duel-channel CNN architecture that trained with high definition images to detect and segment very complicated root systems. This model can segment the plant root system with high accuracy and store resultant root system architecture in RSML format. Below is a summary of the key contributions of our research:

1. To the best of our knowledge, this is the first high definition (HD) root phenotyping system that can segment very complicated and micro details (root hairs) from occluded root systems.
2. A fully automated end-to-end training and infernece architecture that offers complete root segmentation without manual intervention.
3. Developed a new dual-channel CNN architecture that pixel-wise segments whole root system of wheat (primary and small lateral roots).
4. Benchmark results on publicly available wheat roots dataset.
5. Analyzed and compared the proposed CNN model with baseline semantic segmentation networks like FCN, VGG, SegNet, UNet and DeepLab V3.

## II. Related Work

Currently, machine learning offers a lot of capabilities to cover the genotype-phenotype gap. Most state-of-the-art tools and systems provide high-end support to detect and measure plant features. The emergence of deep learning-based image analysis methods has proven to be more suitable and powerful for plant phenotype applications [14]. These applications offer outstanding performance and exceptional results when presented with sufficient amounts of training data [15].

CNN have shown exceptional results in a verity of everyday tasks such as handwriting recognition [16], image classification [17], [18], image segmentation [19], [20], [21] and instance detection and segmentation [22], [23]. Deep learning has recently made its way into plant phenotyping applications due to its reported successes for the detection and segmentation of objects [24]. They allow biologists to phenotype plant traits by feeding raw images from a camera or video stream with very little or no manual intervention. They provide outstanding results for segmenting and or classifying plant traits when compared with the traditional image-based plant phenotyping methods.

Deep learning applied to plant phenotyping problems, including the classification and segmentation of various plant traits [25] and plant disease detection [26], [27]. The recent successes of deep learning in these domains have encouraged researchers to investigate more complicated and challenging problems in plant phenotyping, for example, leaf counting [28], [29], tip detection [30] and lots of other complex tasks [31], [32], [7].

Pound et al. [32] achieved state-of-the-art results by fully automating the feature extraction and localization of wheat plant shoot using deep learning approaches. In another research, Pound et al. [7] developed a comprehensive deep learning model for wheat spikes and spikelets detection using ACID dataset [7]. The proposed models designed to classify and locate the spikes and spikelets with very high accuracy. To bridge the genotype-to-phenotype knowledge gap, Ubbens et al. [29] introduced the idea of deep plant phenomics that offers wide-ranging pre-trained NN models for common plant phenotyping tasks. They release tools that were open-source, offering plant scientists the capability to train models for plant mutant analysis, age regression and leaf counting tasks. Singh et al. [33] developed a state-of-the-art deep learning model for plant stress analysis and offered a comprehensive list of on-going research that uses deep learning approaches for plant stress phenotyping.

Besides the traditional image-based and deep learning plant phenotyping approaches, other machine learning techniques have successfully used to phenotype plant traits [24]. Singh et al. [34], [33] have successfully applied clustering algorithms, support vector machines and neural networks to phenotype various plants traits. Atkinson et al. [15] used Random Forest to assess the architectural features of a quantitative characteristic of plants. They used a semi-automated image analysis method with the Random Forest model to achieve high throughput phenotyping of roots.

Lobet et al. [35] analyzed root systems using a fully automated image analysis pipeline based on the deep learning platform. The work responded to the lack of data for deep learning phenotype traits by developing a synthetic library of images, which improved the detection and estimation of various root traits; however, the key problem with this research is synthetic data. The images used for training are synthetic and not behaving similar to real root. It means lots of real root traits can not be assessed using this system.

The challenges of the traditional image-based phenotyping methods include manual tuning of parameters, limited performance and semi-automated compared with deep learning methods. However, the use of deep learning for plant phenotyping applications is more successful and reports outstanding results [24], [36], [37], [38], [39]. They have the capability of resolving plant phenotypic problems beyond bounds, given the availability of a large amount of data and high-throughput design and helps bridge the genotype-phenotype gap (Yang et al., 2014). They have also been used to solve very complicated phenotypic problems, which were previously not possible with the traditional image-based methods [29], [31].

## III. Methodology

Overview of Proposed Architecture: we have trained a custom-designed CNN architecture for accurate pixel-wise segmentation of wheat root, which we called “RootNet”. A wheat root is sub-divided into primary and secondary (or lateral) roots. Our network detects roots (primary, lateral), seed location and background. The proposed RootNet is a multi-channel CNN architecture with two different pipelines, each dedicated to performing a specific task. We merged the pipelines at the feature fusion unit to form a single channel. We present more details in the sections below:

### A. Network Architecture

In Figure. 1 is the RootNet architecture, which devised of multiple channels to perform efficient segmentation of small size roots features. The architecture made up of a deep and shallow CNN pipeline along with a feature fusion unit.

### B. Motivation

Combining different levels of features is beneficial to the overall performance of the network. For example, PSPNet [40] used an explicit pyramid polling architecture to improve functionality by combining global and local details. ContextNet [41] makes use of a similar approach to feeding networks with different significant features based contexts. Apart from combining different levels of features, deep CNN has been used to improve network performance [42], [43]. They have proved quite efficient in extracting complex and abstract features. However, in the case of roots, especially lateral roots, they appear smaller or invisible when we go deeper into the network. In this case, using a deep CNN is less efficient, especially in detecting lateral roots. We draw our inspiration from ContextNet [41] to make use of a multi-channel and multi-resolution CNN architecture with dual features channels. It is composed of a deep channel Encoder-Decoder (ED) that intended to process high-resolution images and a shallow channel that processes low-resolution images.

### C. CNN Design

This section will describe the overall architecture of the proposed system. RootNet has two channels; one is a deeper Encoder-Decoder branch designed to process the high-resolution image and a shallow branch intended to process a low-resolution image. As we aforementioned, deep networks lose extensive feature details as they go deeper. So, to resolve this issue, we proposed a novel idea to crop images into equal portions and feed the network each portion individually. Each part will be cropped at high resolution and fed to the network for processing. In this way, there is no need to resize the image to a minimal level that leads to the loss of most of the lateral roots features. The full branch will use the (*h × w*) full resolution, while the second branch with lower resolution (e.g. *h/n × w/n*), where *n* = 2^2*n*^. The high-resolution branch will take the cropped high-resolution image in 2^2*n*^ format, which means each high-resolution image will be cropped to given crop value and later used as an individual image for further processing. It is responsible for capturing the tiny details of primary and especially seed location and secondary roots. In contrast, the low-resolution channel is responsible for segmenting global details (e.g. primary root, boundaries). Several experiments have been performed before finalizing the RootNet ultimate design. During these experiments, we have tried different channels, pooling and feature map variations. There are different Encoder-Decoder CNN designs assessed for proposed architecture and the channels mentioned above. We have experimented with FCN [19], VGG [44], SegNet [45], UNet [21], and DeepLab-V3 [46] for assessment of most efficient CNN architecture. The final architectural assessments showed that RootNet dual-channel architecture outperformed the aforementioned CNN systems.

The deep channel is based on an Encoder-Decoder CNN design. In contrast, the shallow channel is a VGG [44] like architecture, where we have embedded 11 convolutions along with Batch Normalization (BN) [47] and Poolings units. In the proposed architecture, it also assessed that the use of pyramid pooling [48] units at the end of the low-resolution channel, offered a great deal of performance boost. Due to a massive difference among class distributions, we make use of the class weight balancing method [49]. For this purpose, we classified the pixel-wise weights of each class and fed it to the network while the loss calculation stage.

### D. Deep ED-CNN Channel

It is hard to train a very high definition image with a very deep CNN architecture, due to current GPU size restrictions. The key purpose of training an HD image is to preserve very small and significant details of features (lateral roots, root hairs). Due to resizing and downsampling images, such essential features get lost. Though, the deep CNN proved to be very useful and offered the highest quality performance, such as ResNet [17] and Inception [50]. So, to extract very import features, we defiantly need a deep CNN channel that can segment our very miniature and complex root topology. Though, we cannot get such details by feeding the low-resolution image to the network. To resolve this issue, we proposed an innovative approach to crop the image into 2^*n*2^ patches and fed our deep CNN network channel. By doing so, we did not compromise our essential roots details.

Each composed patch treated as an individual image. This patch-based approach is a kind of over-sampling method that considers both local patches of the entire image for the final segmentation task. Oversampling is a well-known method for deep CNN to improve network learning capability as it helps to get rid of the overfitting problem without the additional need for data. Motivated by this idea, we crop the given labelled images in *Xi, i* = 1, 2,. *n* into 2^*n*2^ local patches *Xij, j* = 4, 8*m*, where n is the total number of training images and m is the total number of cropped patches belong to each image. The method of cropping is different from traditional cropping methods, which used for data segmentation of the training dataset. As we also use the full image for training on the second channel. This channel emphasis to segment very small details present in the image. Due to the high definition of the image, we preserve lots of important information besides having several pooling and downsampling operations. However, we lose some boundary details, which can be compensated by using a shallow CNN Pipeline.

### E. Shallow CNN Channel

In the multi-channel proposed architecture, the 2nd channel is a shallow channel designed to locate high-level details (primary roots, boundary details). This channel designed to extract global features and strengthen the deep channel segmentation further. It is composed of multiple convolution layers, as architecture shown in Figure. 1. This shallow pipeline takes four times a downsized version of the original image as feed to the deep pipeline. This image propagates through multiple convolutions, activations and pooling to final feature fusion unit. At feature fusion unit all feature map is combined with deep CNN pipeline to construct a comprehensive set of feature maps containing global and local both types of contexts.

### F. Feature Fusion Module

The key idea of the feature fusion unit is to combine two channels into one. RootNet two network channels (deep, shallow) combine their feature map at feature fusion unit. This proposed model combines the high resolution and low-resolution channels to predict the final pixel-wise segmentation of classes. This channel designed to be efficient for reducing the network processing overload. This unit composed of convolution, residual and final segmentation layers. It concatenates all feature maps coming from two channels and train them jointly so the network can learn all missing feature either of two channels. We have used a feature map-based concatenation method that is more efficient than max pooling. There are several reasons to combine both channels into one channel. Neither the patch-based nor the shallow channel alone will be able to predict the final segmentation, as each channel loses several key details during processing. Finally, combining the channels increase the feature maps, thus providing more information for the final pixel-wise segmentation. Another key reason to combine both channels is that the multi-channel pipeline is never perfect and probably biased. Therefore, an image-based fusion model can help to learn and correct the biases among both channels. At this channel, we segment the image into given classes, primary roots, lateral roots, seed location and background. These experiments performed to ensure the right combination layers for feature fusion unit. Finally, it is assessed that placing a residual block with 1×1 convolution can offer outstanding results. Further details of network architecture and each unit are presented in Figure.1. In Eq. 1 we have outlined the features fusion method:

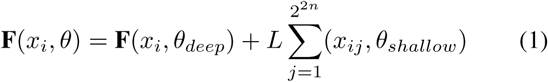

Where, **F**(*x*_*i*_, *θ*) is overall final set of feature maps processed by feature fusion unit. The **F**(*x*_*i*_, *θ*_*deep*_) depicts the deep and 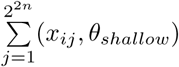 shows the shallow channels of CNN architecture. Where *L* is a balancing factor, as it is used to balance the effect of two different networks.

## IV. Experiment

In this section, we present the details of a quantitative study performed to ensure the performance of the RootNet CNN system. The results produced are performance benchmark on Bread wheat (Triticum aestivum L.) dataset [7] for semantic segmentation. Figure 3 shows the overall workflow chart of the proposed experiment. Further details about experiment setup and dataset outlined in the section below:

**Figure 2:**
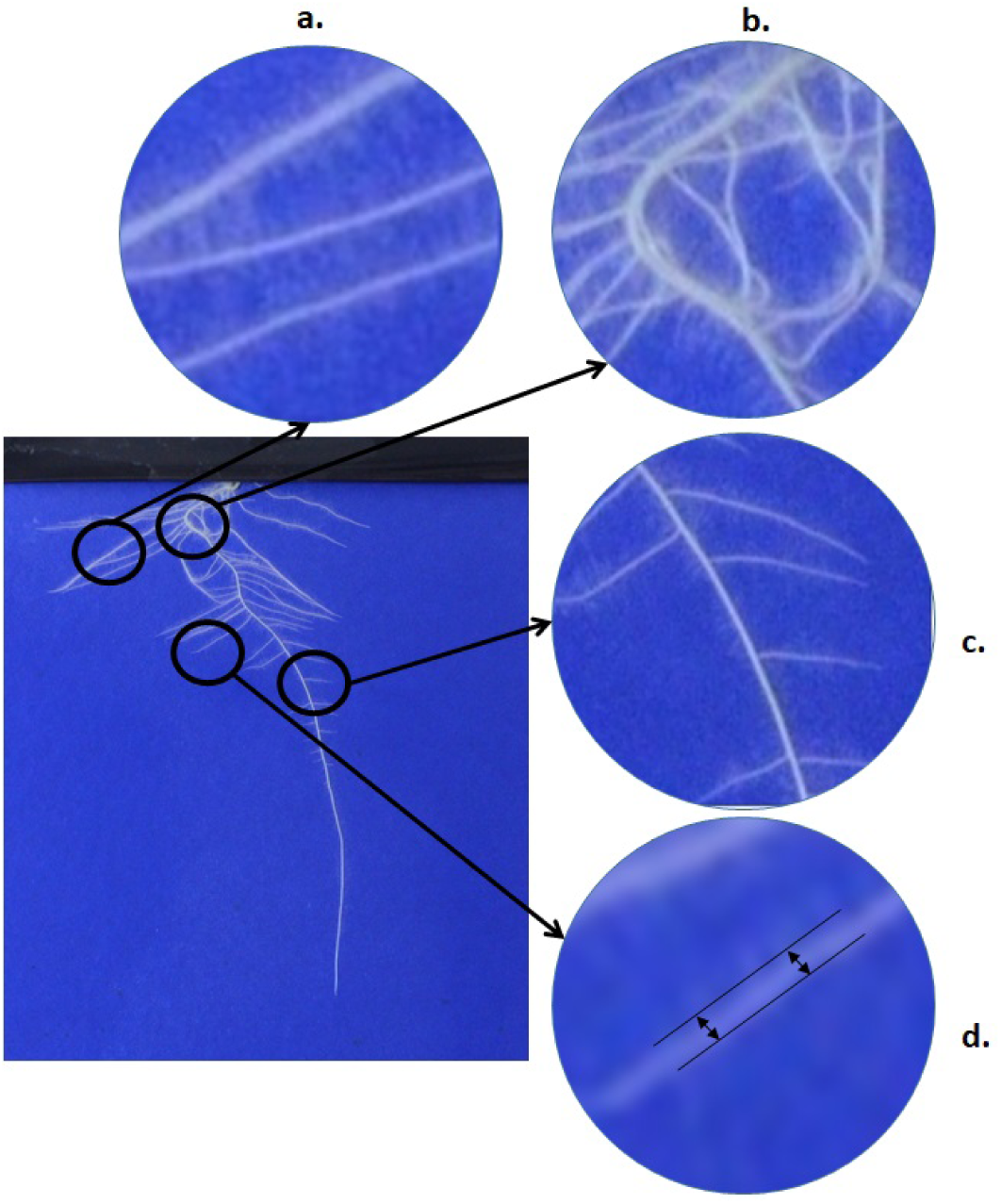
A sample image from the dataset of wheat (Triticum aestivum L.). a) An example of the challenging root intersection that makes harder to count and segment individual roots. b) It is challenging to extract root topology from complex occlusion in natural root growth using traditional or machine learning-based methods.c) There is a set of lateral roots that are very difficult to annotate, even using manual annotation methods. d) Very small root hairs (1-3 pixel wide) leads to classification difficult and sometimes goes undetected when imaged resized to lower dimensions.

**Figure 3:**
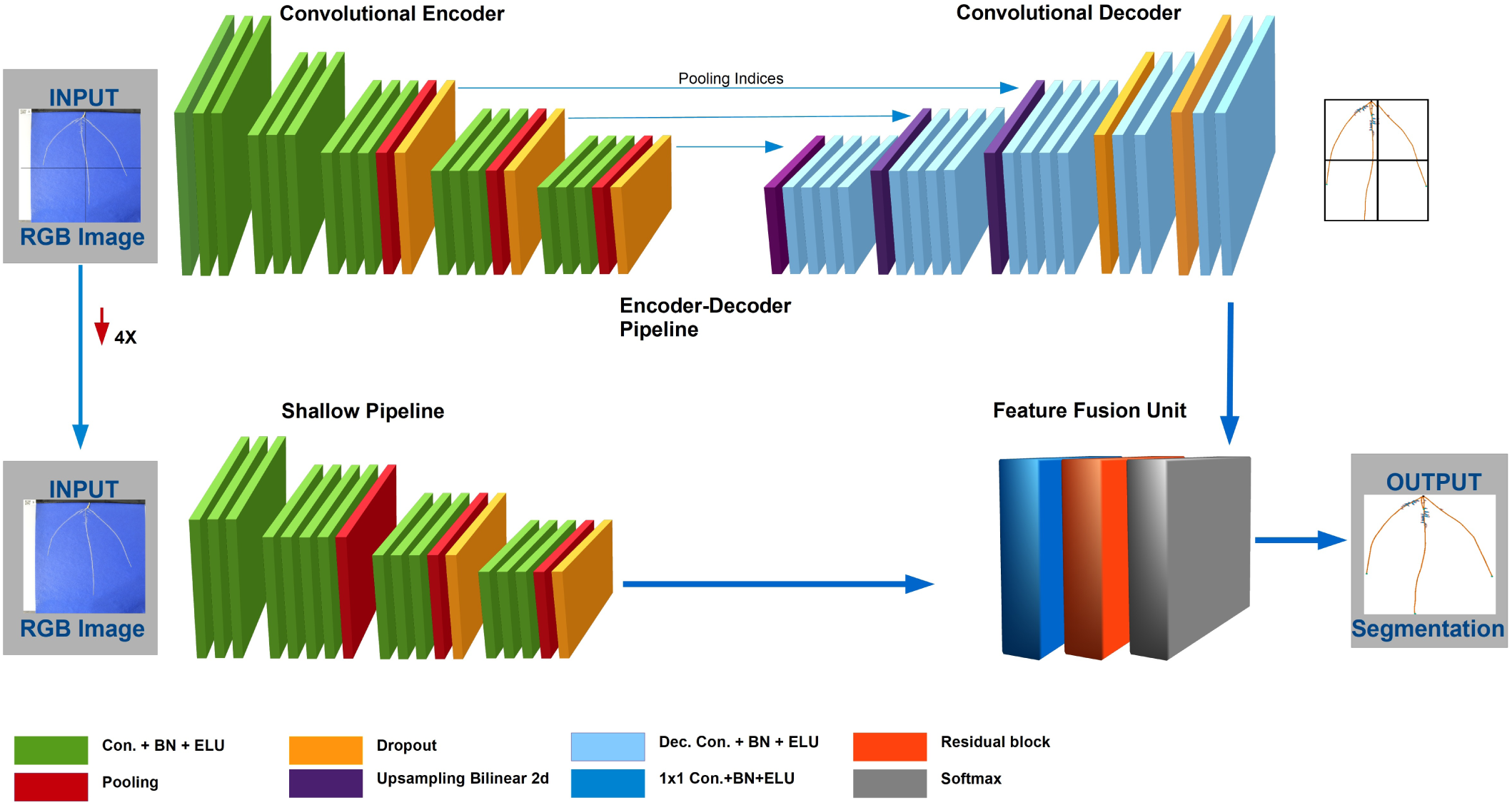
Layers Architecture of RootNet CNN

### A. Implementation details

The overall RootNet architecture designed using the Pytorch [51] deep learning library. We performed experiments on two Nvidia Titan X GPUs with Cuda 8.0 and CuDNN v6 support. We used exponential linear unit (ELU) [52] as non-linearity function with Batch Normalization (BN) at all layers. For better network convergence, we train each channel individually and later combined to re-train (using transfer learning method) for effective segmentation. We trained RootNet using RMS-Prop optimizer [53] with a weight decay of 0.9 and a momentum of 0.9. We then trained the network for 500 epochs with constant updates in the learning rate. The initial learning rate set to 0.1, which reduced by factors of 10 every 50 epochs. The results are outlined using overall mean accuracy and mean intersection over union (MIoU). To assess the accuracy of the system is a much better way we make use of two famous accuracy assessment metrics Global Accuracy (GA) and Class Average Accuracy (CAA). The GA offers an overall percentage of pixels properly classified in the dataset, while CAA used to assess the predictive accuracy mean of the entire classes.

### B. Wheat Root Dataset

The bread wheat (Triticum aestivum L.) [7], [54] dataset composed of a total of 3682 high-resolution images with 2570×1900 size each. 2946 images used for training and 736 images for the validation dataset. These roots are grown in viable pouches in a controlled environment. Parameters for roots growth set with 12 hours photoperiod with daytime temperatures of 20 degrees and nighttime temperatures of 15 degrees the light intensity set to 400 umol m25-1 PAR as per (atkinoson 14). The images are taken after 9-day growth using the Nikon D5100 camera. The root system architecture extracted using RootNav software [13]. We extracted ground truth images of the dataset using the Root System Markup Language (RSML) [55] files that offer root architectural details and growth geometry.

### C. Dual-Channels CNN

The design of RootNet finalized after multiple number of experiments. The number of channels was limited to two branches. Earlier research shows that increasing the depth of the network increases the overall accuracy of the network. However, this presents a negative impact on the network resources consumption: longer training time and more GPU memory. Therefore, designing a single, very deep channel can introduce similar issues to the network. The key idea to keep the number of channels to two is to reduce the network depth-size and reduce overall resource consumption. Two channels with less depth than one single super-deep network can offer better performance with low resource consumption.

### D. Input Channel Resolution

The available dataset offers an opportunity for different input size resolution with each network branch. The image resolution in our dataset is 2500×1900 pixels, which cannot be feed to the network directly due to GPUs size restriction. Therefore, we have to crop the images. For experimental purposes, we found that 1024×1024 pixels size works for our higher resolution branch and 256×256 to work for the low-resolution branch. Later, each cropped image is put together and fed to a feature fusion unit, where it combined with the low-resolution channel for predicting the output. The feature fusion unit makes use of combined features to predict the final pixel-wise segmentation map of the input image.

### E. Results and Analysis

RootNet has trained 500 epochs and tested for bread wheat roots dataset. For analysis of performance, we have experimented with number CNN architectures, including FCN8, VGG16, SegNet, Unet and DeepLab-V3. During the training of these networks, we realized that these networks struggle to converge due to the microscopic size of roots features. However, the proposed approach offers capabilities to detect minuscule details inside root system architecture. As a result, we get a very higher-class classification rate. Figure. 4 shows a comparison of input, GT and output image, which shows that proposed CNN architecture performs very efficiently in terms of segmenting each class separately and detecting more roots in comparisons to give GT image.

**Figure 4:**
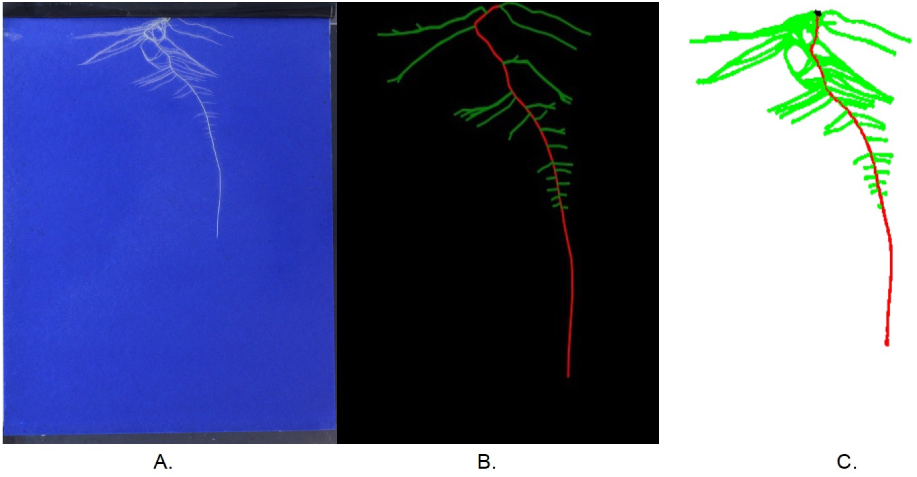
Output segmentation results analysis. **(A)** Input image. **(B)** Ground Truth extracted from RSML file (anotated by biologists). **(C)** Segmentation Results of RSML

The Figure. 5 shows RootNet final results in graphical form. The graph Figure. 5A shows the test error versus the training error of the proposed system. The figure shows the Training/Test Loss curve of proposed CNN. It can be seen that test loss reduce down gradually and start stabilizing after reaching its minimum level. Our method of constant learning rate reduction helps us to improve the network accuracy and reduce loss efficiently. The Figure. 5 graph shows the MIoU of RootNet during the validation process. We have used mIoU metric as it is a more stringent metric as compared to standard class average accuracy as it penalizes false positives and it is also the most well know metric for benchmarking network performance. Here we can see a great deal of enhanced performance in RootNet resultant scores as other networks are struggling with the segmentation of crucial root features.

**Figure 5:**
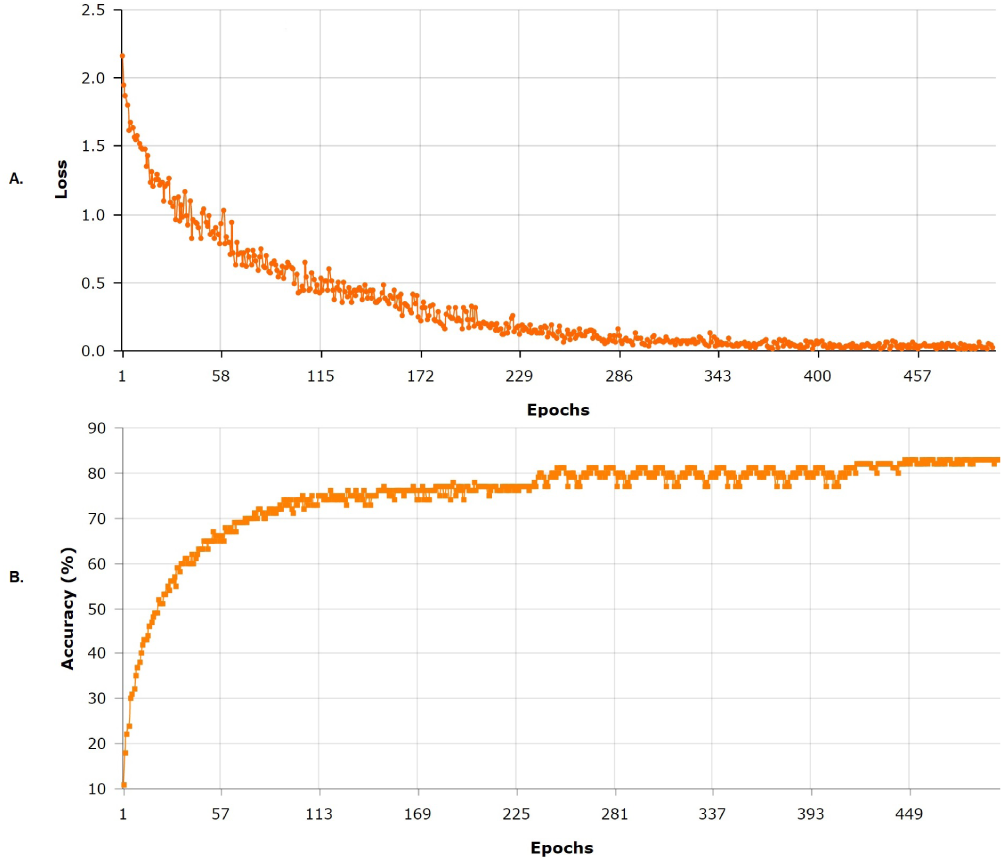
Performance Analysis. **(A)** Comparison of Test Error v/s Training Error. **(B)** Mean Intersection over Union (MIoU) on validation set

Table. I shows the run times of different resolutions where RootNet achieves very high accuracy at offering a very higher performance in terms of the inference time of the network. The key purpose of this analysis to benchmark the performance of the proposed network for semantic segmentation. The experiments carried out by keeping all hyper-parameters at the same level. The results are shown in Table. I. It observed that the proposed CNN architecture outperforms all CNNs mentioned above architectures and has the highest performance. The test loss, AC, ACC and IoU used as two main performance assessment parameters. In Appendix section the Figure. 6 shows a graphical results analysis of RootNet with the aforementioned CNN architectures. It can be seen that RootNet is successfully able to segment and assess almost every detail in the given image.

**Table I:** Quantitative Comparison: This table shows a quantitative comparison of RootNet with well-know earlier CNN architectures for roots segmentation problem. RootNet outperforms all the other systems. We have used global average, class average and mIoU as standard metrics for comparison of segmentation results.

| CNN Arch. | Global avg. | Class avg. | mIoU |
| --- | --- | --- | --- |
| VGG [44] | 37.32 | 38.5 | 31.11 |
| FCN [19] | 43.78 | 48.56 | 39.47 |
| SegNet [45] | 68.68 | 70.23 | 47.05 |
| UNet [21] | 70.47 | 69.65 | 51.88 |
| DeepLab-V3 [46] | 87.70 | 85.90 | 54.0 |
| **RootNet** | **96.4** | **91.35** | **62.19** |

**Figure 6:**
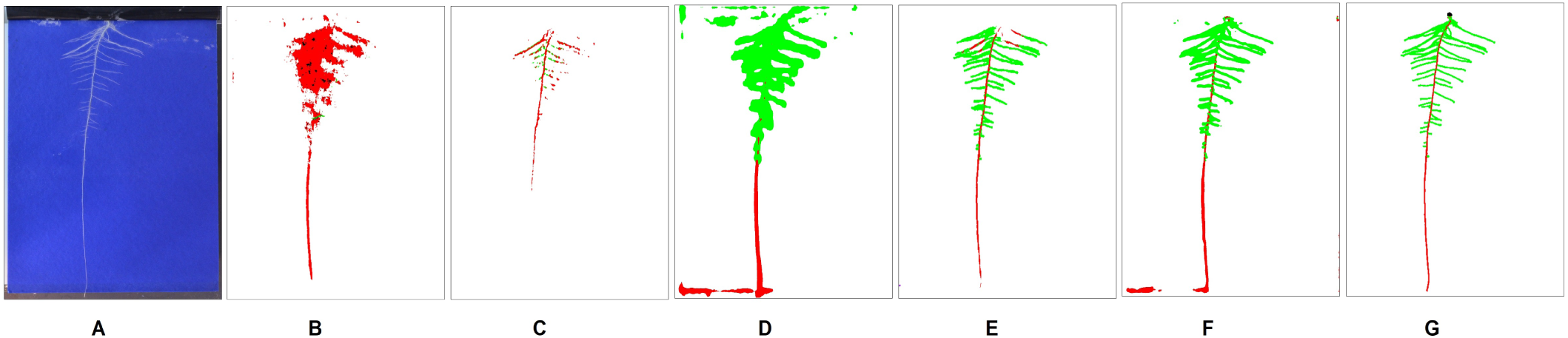
Segmentation Results Analysis. This figure provides a visual analysis of segmentation results extracted from all above mentioned CNN architectures. We have used one of the most challenging images to assess the performance of the proposed network. This image contains a cluttered background and a large number of microscopic lateral roots. **A** Input Image **B** VGG [44]. **C** FCN [19]. **D** SegNet [45]. **E** UNet [21]. **F** DeepLab-V3 [46]. **G** RootNet.

## V. Conclusion

RootNet aimed to offer better segmentation without sacrificing pixel details to maximise detection accuracy ultimately. The proposed reserch has shown a great deal of sucess in terms of segmenting HD root images to extract maximum details from occluded roots structures. The proposed CNN architecture offered a great deal of capabilities to distinguish among different levels of roots (primary, lateral). The final results have shown that RootNet achieves very high classfication accuracy and performance in terms of per-pixel segmentation of the root system architecture. The fundamental purpose of this analysis to benchmark the performance of the proposed network for root systems semantic segmentation. The experiments carried out (with different CNN architectures) by keeping all hyperparameters at the same level. It observed that the proposed CNN architecture outperforms all CNNs mentioned and has the highest performance.

## VI. Future work

For future work, we aim to extend the current research to other plant species. The intention is to design a wideranging single CNN architecture suitable for root detection and segmentation for a wide range of plant species. We will also intend to detect more root features (e.g., root tips). To further extended the scope of research for more complex operations (e.g., count roots, measure lengths and diameter, virtual reconstruction).

## VII. APPENDIX

### Appendix A: Bibliometric Analysis

Plant phenotyping emerged as an extensive research area as a result of widespread advancements in image processing and machine learning technologies. It is the quantitive description of plants annotation, ontologies, physiological and biochemical properties [4]. There are two broad categories that plant phenotyping seeks to study; the first one is the above-ground and the second one is below-ground. The above-ground phenotyping study the quantitative measurement of the plant shoot and the below-ground studies the root system of the plant. Whereas, the above-ground phenotyping has extensively studied; however, the below-ground phenotyping is an emerging area with very few state-of-the-art image-based tools. The challenge with below-ground phenotyping is that it is difficult to use a simple imaging device to capture the whole root system without damaging or disturbing the natural growth process of the plant, thus making it difficult to acquire lots of data for their study. Even if there are data readily available, there is another challenge that usually the roots are very tiny or arranged in a complicated arrangement; therefore, image-based approaches prove difficult or sometimes impossible to segment the whole root from the soil. Machine learning (ML) helps in variety of biological challenges such as genome annotations [60], gene function prediction [59], genetic variation predictions [58], genetic region identifications [15] and most importantly image-based plant phenotyping [57]. Image-based plant phenotyping methods topped over traditional ML-based techniques [57]. ML-enabled researchers to capture and accurately predict plant phenotypes with a less amount of time and effort as compared to manual ML analysis. Earlier ML technologies and methods are mostly data-driven with tools to assess and analyze roots morphological parameters [1]. With gradual growth in image acquisition and analysis methods, it becomes possible to apply image-based plant phenotyping [56]. A large image dataset helps to train the system to assess roots growth and anatomy. Recently, methods and tools have emerged for image-based root phenotyping [4], [6], [5], [7]. However, these tools are partially automated and require some manual intervention to perform specific phenotyping analysis. There are several fully automated tools [62], [39], [61], which are available for the above-ground plant phenotyping analysis but these tools are not equally applicable to below-ground plant phenotyping analysis (e.g., roots cross-section).

To analyze the emergence of computer vision and deep learning use in plant phenotyping, we have performed a literature survey. For this purpose, we have extracted research literature from two top literature resources Web of Science and Scopus. We have extracted research publications (article, conference papers, reviews, books, chapter) from the year 2010 to 2019. We searched research literature for plant phenotyping and the use of technology for phenotyping. After careful analysis and elimination of irrelevant records to we have extracted 1556 research publications for ranging on a span of more than nine years. Later we have used literature analysis tools BibExcel and VOSviewer to filter and visualize technology trends in plant phenotyping paradigm. In the Figure. 7 offers an overlay visualization of research literature from 2010 to 2019. In this overlay visualization graph, different colors used to demonstrate the various technology trends each year related to plant phenotyping. We can observe an emerging trend of using machine learning and deep learning for plant phenotyping since 2017.

**Figure 7:**
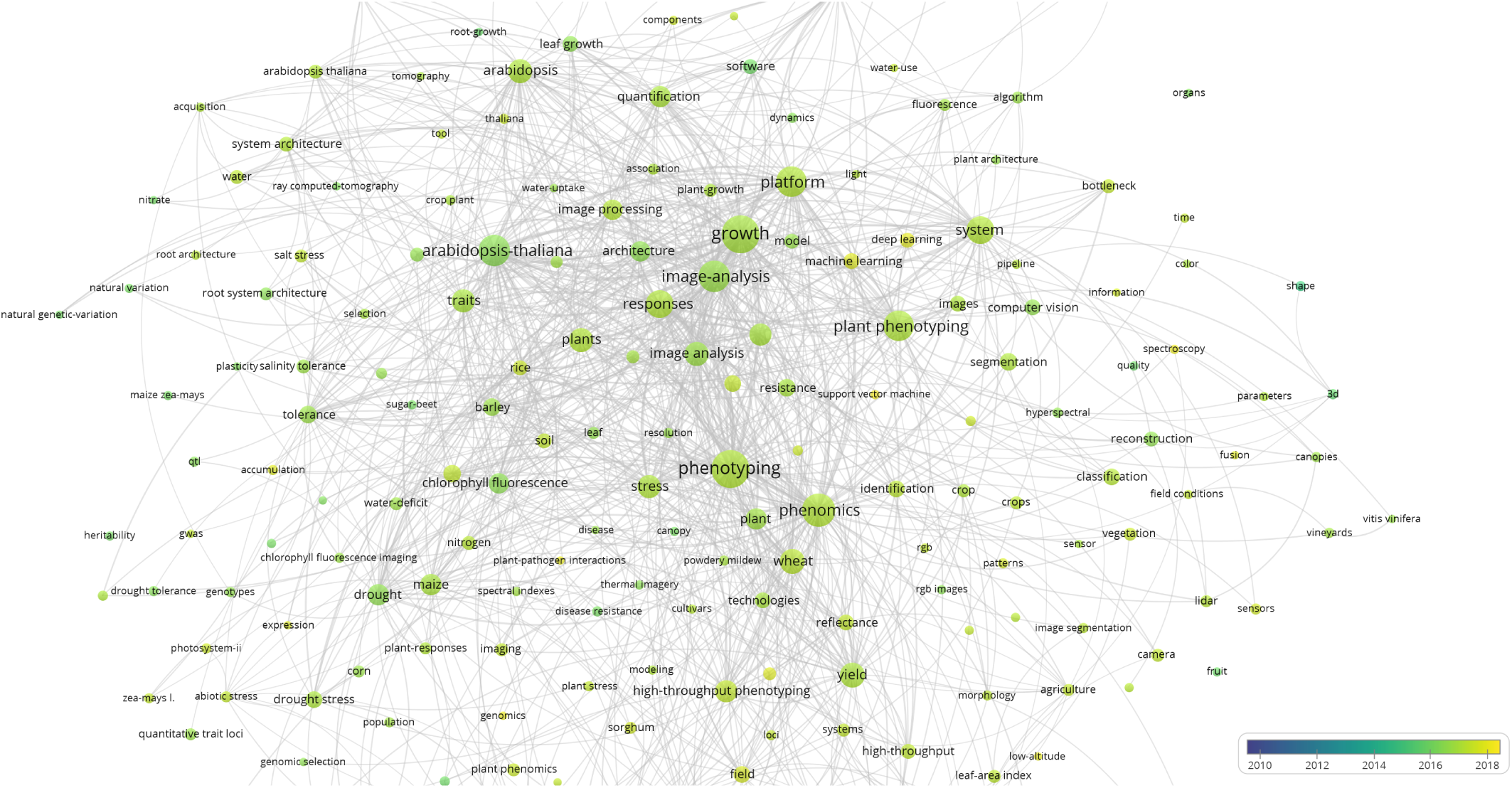
Overlay Graph of Plant Phenotyping Literature 2010-2019

To assess the root phenotyping trends over the years, we have drawn the Density Visualization graph in Figure 8. This graph is based of same above mentioned literature data. This graph shows a significant research gap between mainstream plant phenotyping and root phenotyping. In this graph, the red cluster is the most highly published research domain, while the yellow and green clusters are less mainstream plant phenotyping research topics. Root segmentation, root analysis, and overall root phenotyping is one of the most neglected research domain. Root phenotyping almost lays out the plant-phenotyping research domain clusters. This is one of the fundamental reasons and motivations behind this research to make root phenotyping as significant as above-ground plant phenotyping.

**Figure 8:**
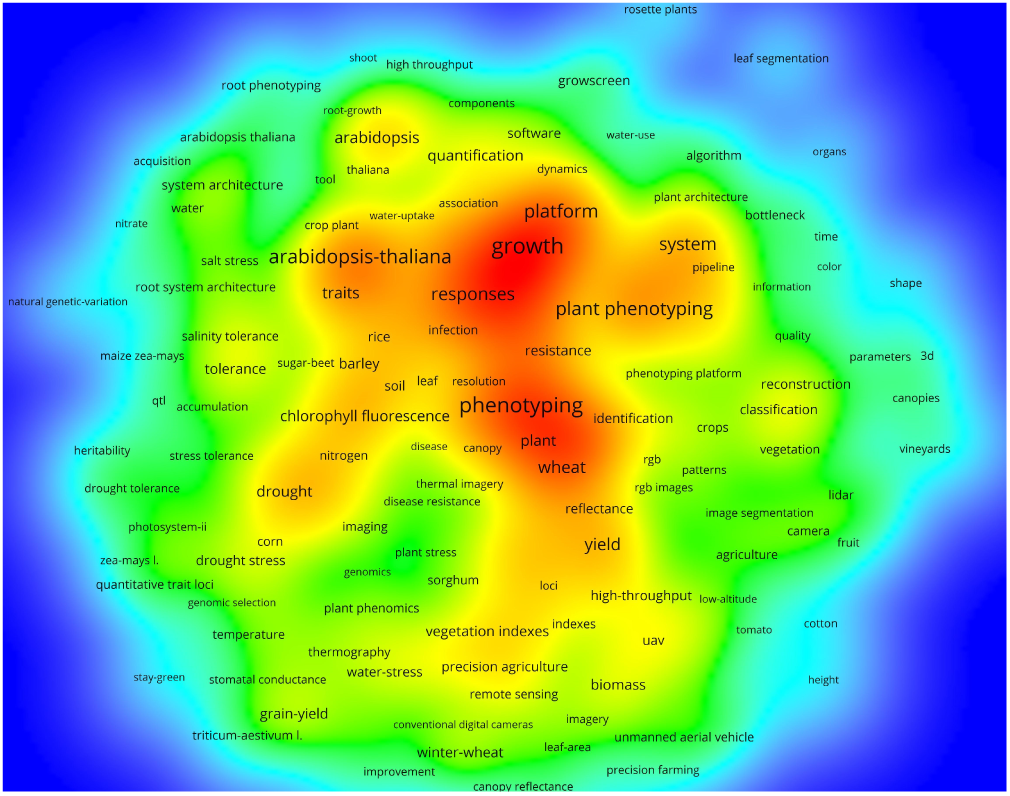
Density Visualization graph of Plant Phenotyping Literature 2010-2019

Root phenotyping is one of the critical research areas that demands a great deal of attention to offer more better plants analysis and support future food safety research. However, the use of machine learning and technology-based systems is currently very limited in this domain. We have extracted a small fraction of the literature, as mentioned earlier, network visualization graph in Figure 9. This small fraction highlights the earlier and current trends in root phenotyping research. None of the above-shown patterns are related to the use of machine learning or deep learning technologies. The use of new neural network-based analysis is an emerging trend in root phenotyping that holds a great deal of potential to unleash the complicated features of roots and help biologists to under roots a much more better way. To deal with the above-mentioned root phenotyping research gap, we proposed RootNet a fully automated end to end trained CNN system that pixel-wise segments whole wheat roots.

**Figure 9:**
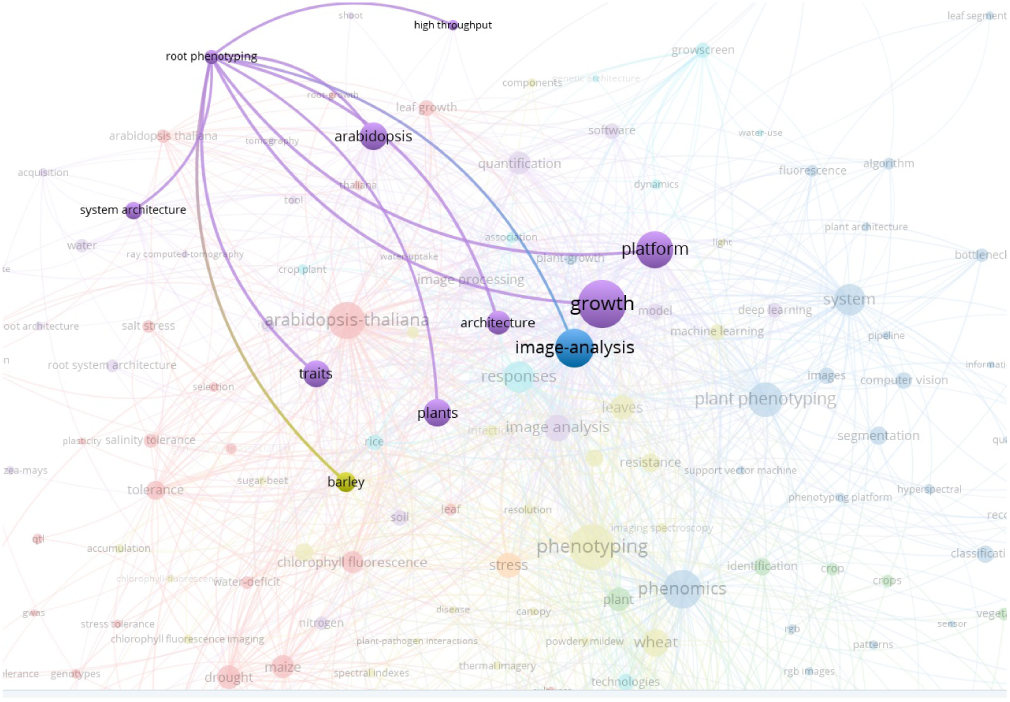
Network Visualization graph of Plant Phenotyping Literature 2010-2019

### Appendix B: Additional Segmentation Results

The Figure 10 shows an additional set of dataset samples segmentation results.

**Figure 10:**
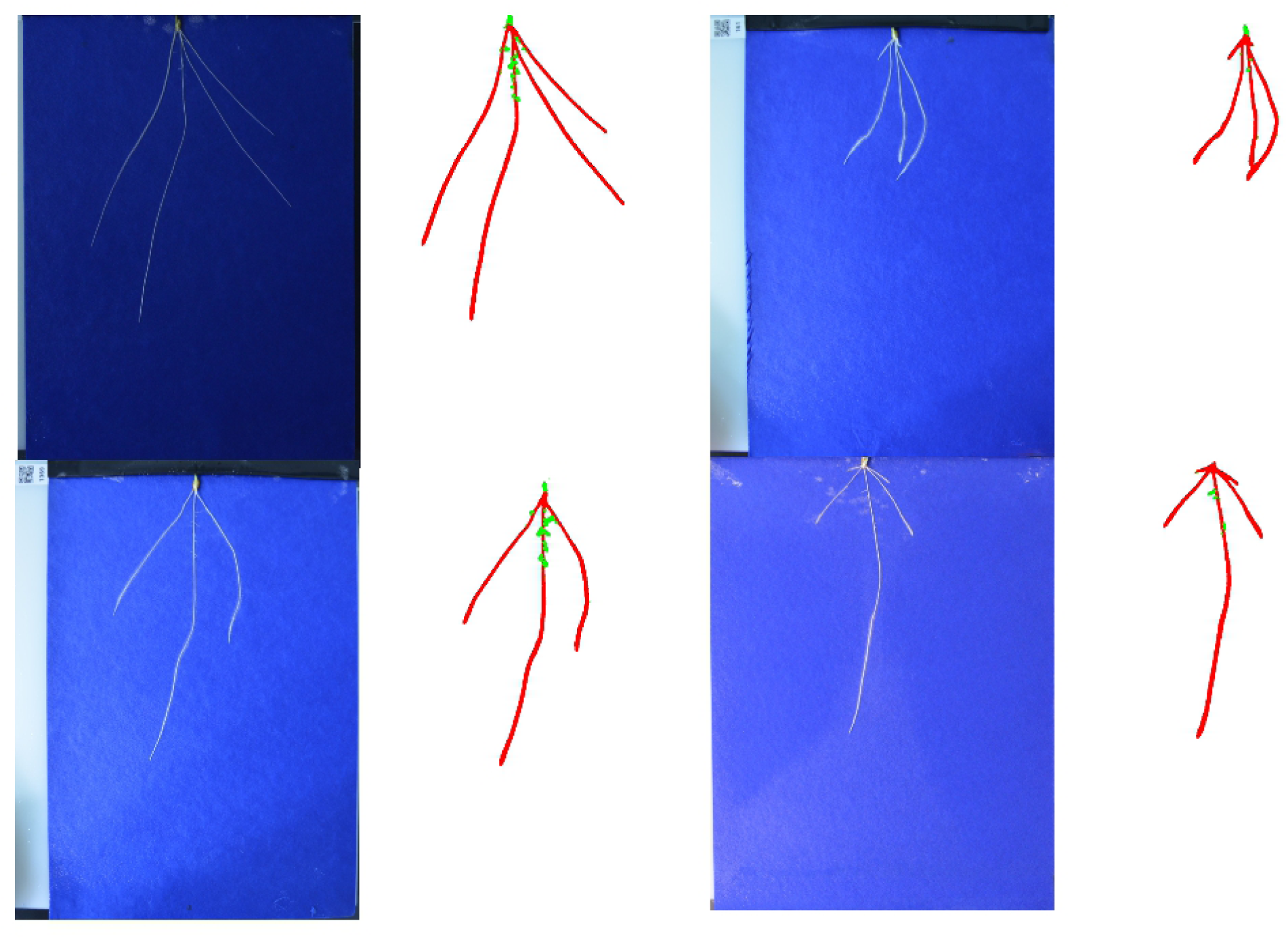
Additional dataset samples for Segmentation Results Analysis.

